# Genomic signatures of extensive inbreeding in Isle Royale wolves, a population on the threshold of extinction

**DOI:** 10.1101/440511

**Authors:** Jacqueline A. Robinson, Jannikke Räikkönen, Leah M. Vucetich, John A. Vucetich, Rolf O. Peterson, Kirk E. Lohmueller, Robert K. Wayne

## Abstract

The observation that small, isolated populations often suffer reduced fitness as a result of inbreeding depression has guided conservation theory and practice for decades. However, investigating the genome-wide dynamics associated with inbreeding depression in natural populations is only now feasible with relatively inexpensive sequencing technology and annotated reference genomes. To characterize the genome-wide effects of intense inbreeding and isolation, we sequenced complete genomes from an iconic inbred population, the gray wolves (*Canis lupus*) of Isle Royale. Through comparison with other wolf genomes from a variety of demographic histories, we found that Isle Royale wolf genomes contain extensive runs of homozygosity, but neither the overall level of heterozygosity nor the number of deleterious variants per genome were reliable predictors of inbreeding depression. These findings are consistent with the hypothesis that severe inbreeding depression results from increased homozygosity of strongly deleterious recessive mutations, which are more prevalent in historically large source populations. Our results have particular relevance in light of the recently proposed reintroduction of wolves to Isle Royale, as well as broader implications for management of genetic variation in the fragmented landscape of the modern world.

## Introduction

Under increasing human population pressure, many species with once continuous ranges have been reduced to small, fragmented populations (Ceballos and Ehrlich 2014). Higher levels of inbreeding in such small populations may elevate the risk of extinction through inbreeding depression. Large carnivores are particularly susceptible, since they typically have lower population densities relative to the herbivores they prey on, and they often require extensive natural areas to persist. Further, they are frequently persecuted because of real or perceived threats to humans and livestock (Ripple et al. 2014). Some well-known examples of inbreeding depression in the wild have been observed in large carnivores, such as Florida panthers (*Puma concolor*, Roelke et al. 1993) and gray wolves (Liberg et al. 2005; Räikkönen et al. 2006, 2009). Studies of inbred wolves in the wild and in captivity have found elevated rates of blindness, cryptorchidism, heart and kidney defects, dental anomalies, and vertebral malformations, as well as decreased reproduction, litter size, body weight, and longevity (Laikre and Ryman 1991; Laikre et al. 1993; Liberg et al. 2005; Fredrickson et al. 2007; Räikkönen et al. 2013; Åkesson et al. 2016). Phenotypic defects associated with inbreeding are not limited to carnivores, however, and have been observed in numerous plant and animal species (Keller and Waller 2002). Quantifying and maintaining genetic diversity to minimize the risk of inbreeding depression therefore remains a fundamental goal of conservation biology.

Although it has been studied for more than a century, the underlying genetic basis of inbreeding depression remains unclear. Previous studies largely support the hypothesis that inbreeding leads to increased homozygosity of strongly deleterious recessive alleles, which are hidden from selection by remaining in the heterozygous state in large outbreeding populations (reviewed in Charlesworth and Willis 2009). However, the adverse consequences of small population size have been debated, in part because theory predicts that smaller populations may actually have an enhanced capacity for purging strongly deleterious recessive mutations (Hedrick and Garcia-Dorado 2016). With the ever-decreasing costs of whole genome sequencing, it is now feasible to estimate the genome-wide burden of deleterious variants (genetic load) (eg. Lohmueller et al. 2008; Renaut and Rieseberg 2015; Marsden et al. 2016). However, recent studies have primarily dealt with the effects of long-term reduced population size or ancient bottlenecks, such as in non-African human populations or in domesticated species, rather than inbreeding in small populations. Additionally, studies have focused on the excess of deleterious variants associated with expanding populations (Peischl and Excoffier 2015). Generally, small increases in the number of derived deleterious variants per genome (additive genetic load) due to ancient bottlenecks or long-term reduced effective population size have been observed, but the genomic effects of severe inbreeding may have a distinct impact on patterns of deleterious variation.

In this study, we present results from complete genome sequencing of a small highly inbred population of wolves on Isle Royale in Lake Superior that has been under annual observation almost since its founding, serving as a model system for the study of ecological and behavioral dynamics, as well as conservation genetics for decades (eg. Allen and Mech 1963; Wayne et al. 1991; Peterson et al. 2014). The island was likely first colonized by two to three wolves that crossed frozen Lake Superior from the mainland in the 1940s, establishing a population on Isle Royale that, at its peak, included approximately 50 individuals (Peterson et al. 2014). Following a disease outbreak in the early 1980s, the population crashed to 14 individuals and failed to rebound for approximately 15 years. The population exhibited a significant but short-lived improvement in numbers as a response to genetic rescue after a single wolf migrated from the mainland in 1997, before falling to even lower numbers by 2010 (Adams et al. 2011; Hedrick et al. 2014; Peterson et al. 2014). The population is now extremely inbred, and has continued to wane while exhibiting signs of severe inbreeding depression (Räikkönen et al. 2009; Hedrick et al. 2014). A pair of wolves, both father-daughter and half-siblings, descended from a legacy of repeated close inbreeding events, were the only individuals that remained by early 2018, by which time they were either in or approaching senescence. No successful reproduction has occurred since 2014 (Peterson and Vucetich 2017), and the population is expected to disappear without the reintroduction of wolves by humans, a move that is under review by the National Park Service. Although previous investigation of Isle Royale wolves focused on inbreeding using genetic assays (Wayne et al. 1991; Adams et al. 2011; Hedrick et al. 2014), or morphological assessment (Räikkönen et al. 2009), this is the first study to combine both approaches, and to use complete genome sequence data.

In the past, levels of diversity at relatively few loci have been used to make inferences about the genetic health of populations. These methods may be misleading, because the relationship between heterozygosity and inbreeding depression is not straightforward, but depends on the nature of segregating deleterious variation, which is influenced by demographic history. Our analysis of complete genomes supports the importance of the genomic landscape in assessments of inbreeding. We find that inbreeding depression is likely caused by increased homozygosity of strongly deleterious recessive mutations due to recent inbreeding, rather than an overall increase in the burden of deleterious alleles due to long-term small population size. These results have implications for understanding the genetic basis of inbreeding depression and for the effective management of small isolated populations, particularly those recently derived from large outbred populations, as is the case for many species threatened by habitat fragmentation and loss due to recent human impacts on the landscape.

## Results

### Genomic data set

We obtained genetic samples from eleven Isle Royale wolves collected between 1988 and 2012 for whole genome sequencing and analysis (Table S1). The population size on the island was estimated to number 8-30 individuals during this period (Peterson and Vucetich 2017). A pedigree, adapted from Hedrick et al. (2014), shows the relationships of the sequenced wolves (where known), as well as inbreeding events between close relatives (Fig. 1A). Based on this pedigree, the wolves in our dataset include inbred and putatively non-inbred individuals, with inbreeding coefficients ranging from 0 to 0.375.

**Fig. 1.**
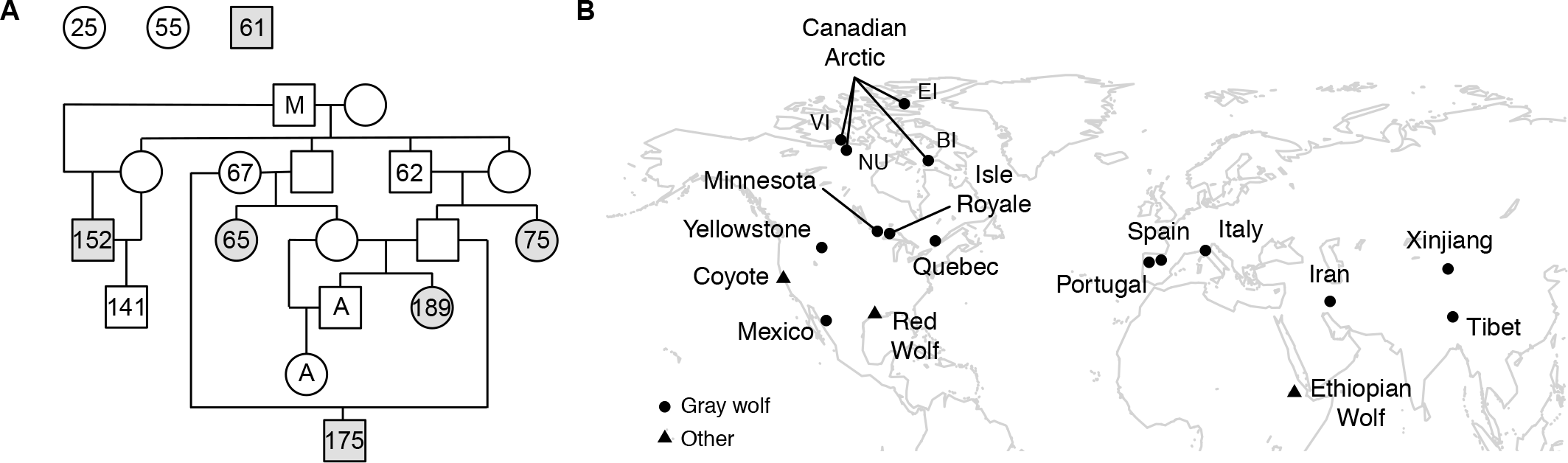
Isle Royale wolf pedigree and geographic origins of genomes in this study. (**A**) Pedigree of Isle Royale wolves sequenced in this study (numbered individuals), adapted from Hedrick et al. (2014). Circles represent females and squares represent males. Relationships were inferred from genotypes at 18 microsatellite loci. Shaded individuals were examined for the presence of vertebral abnormalities for this study (see Table 1). The ancestries of F25, F55, M61, and F67 are unknown. “M” was a wolf that migrated from the mainland in 1997 (Adams et al. 2011). The “A” individuals are the last two wolves alive on Isle Royale as of 2018. (**B**) Map showing approximate origins of all individuals analyzed in this study. Abbreviations: BI, Baffin Island; EI, Ellesmere Island; NU, Nunavut; VI, Victoria Island. See Table S1 for further sample information.

We supplemented the Isle Royale wolf genome sequences with publicly available and newly generated genomes from other wolves and related species (Fig. 1B, Table S1). In addition to the Isle Royale wolves, our complete dataset includes six mainland wolves from nearby Minnesota, nine gray wolves from elsewhere in North America, six Eurasian gray wolves, and a single genome from each of the following species: red wolf (*C. rufus*), coyote (*C. latrans*), and Ethiopian wolf (*C. simensis*). All genomes were aligned, genotyped, and annotated with respect to the domestic dog reference genome (canFam3.1), yielding mean genome-wide coverage values of 9-49X after read filtering (Table S1). We polarized alleles as ancestral or derived using genomes from an African golden wolf (*C. anthus*) and a gray fox (*Urocyon cinereoargenteus*). A cladogram showing the phylogenetic relationships between wolf populations and sister taxa is shown in Fig. S1. Our dataset spans the Holarctic range of the gray wolf and contains individuals derived from a variety of demographic histories that feature recent inbreeding, long-term small population size, isolation, and admixture (Table S2). To our knowledge, Isle Royale and Mexican wolves are the only populations we sampled that suffer from documented inbreeding depression (Fredrickson et al. 2007; Räikkönen et al. 2009).

**Table 1.** Vertebral phenotypes of Isle Royale wolves. Six individuals were examined for vertebral defects, and their phenotypes are listed. LSTV: lumbosacral transitional vertebra, SCTV: sacrococcygeal transitional vertebra, TLTV: thoracolumbar transitional vertebra. “N/A” indicates that the bones from this individual could not be recovered and therefore were not examined.

| ID | Vertebrae examined | Phenotype | Other observations |
| --- | --- | --- | --- |
| F25 | N/A | N/A | N/A |
| F55 | N/A | N/A | N/A |
| M61 | Atlas-Sacrum | No abnormalities | N/A |
| M62 | N/A | N/A | N/A |
| F65 | Atlas-T12, L1-Co3 | Minor phenotypic LSTV | One thoracic vertebra missing from collection |
| F67 | N/A | N/A | N/A |
| F75 | Atlas-Co16 | LSTV, TLTV, SCTV, 2 extra ribs, osteophytes | All of F75 pups had an extra presacral vertebra, 7 of the 8 pups had extra ribs |
| M141 | N/A | N/A | N/A |
| M152 | Atlas-Sacrum | TLTV, SCTV, extra rib, osteophytes | N/A |
| M175 | Atlas-Co9 | LSTV, TLTV, SCTV, asymmetry at Co2, osteophytes | Some ribs are missing from collection |
| F189 | Atlas-Co2 | TLTV, extra vertebra, SCTV, cervical intrasegmental transitional & asymmetry | Some ribs are missing from collection. The cervical malformation is the same type observed in specimen 3529 (see Fig. 3 of Raikonen et al. 2009). |

### Phenotypic evidence of inbreeding depression in Isle Royale wolves

The wolf spinal column is composed of seven cervical (C1–7), thirteen thoracic (T1–13), seven lumbar (L1–7), and three fused sacral (S1–3) vertebrae from atlas to sacrum, plus ~20 coccygeal vertebrae (Co1–22) in the tail. A 2009 study found a high prevalence of vertebral anomalies in Isle Royale wolves sampled between 1964 and 2007, including extra vertebrae, and defects such as thoracolumbar, lumbosacral, and sacrococcygeal transitional vertebrae (exhibiting characteristics of two different types of vertebrae), as well as vertebrae with severe asymmetries (Räikkönen et al. 2009). We examined skeletal remains collected post-mortem from 6 of the 11 wolves sequenced for this study, as well as a litter of 8 pups, to identify congenital malformations, specifically in the spine and rib cage. Only one of these specimens was part of the 2009 study (F65). The incidence of observed anatomical defects was high. One individual was free of aberrations (M61) and one (F65) had a minor transitional variation, whereas the remaining individuals (F75, M152, M175, F189) possessed 2-4 abnormalities each, including transitional vertebrae, extra vertebrae, and extra ribs (Table 1). One wolf, F75, was the product of a brother-sister mating, and had three transitional vertebrae and an extra pair of ribs. This wolf died while giving birth to a litter of pups thought to have been sired by her father, M62 (Hedrick et al. 2014). The pups died within hours, and our examination revealed that all eight had extra vertebrae, and all but one had extra ribs.

The fitness effects of the vertebral defects observed in Isle Royale wolves are not clear, but lumbosacral transitional vertebrae (LSTV) in dogs are reportedly a predisposing factor of cauda equina syndrome, which can cause severe pain, incontinence, gait problems and paralysis (Morgan et al. 1993; Flückiger et al. 2006). LSTV are also reportedly related to asymmetrical hip joint development and secondary osteoarthritis (Flückiger et al. 2017). There are many phenotypic variations of LSTV (Morgan et al. 1993; Damur-Djuric et al. 2006); wolf F65 exhibited only a minor aberration that presumably would have no impact on fitness. Two other wolves with LSTV (F75, M175) showed severe changes with a combination of malformed features. Rarely observed in large, outbred wolf populations, LSTV were previously shown to have steadily increased in prevalence in Isle Royale wolves between 1964 and 2007, a period in which the Isle Royale wolf population became increasingly inbred (Räikkönen et al. 2009). The incidence of LSTV in outbred wolf populations in Finland and historic Scandinavia is 0-1%, compared to 10% in modern inbred Scandinavian wolves, and 33% in Isle Royale wolves prior to 2007 (Räikkönen et al. 2006, 2009).

Additionally, three of the six specimens (F75, M152, M175) also exhibited spondylosis deformans, a condition in which the vertebrae possess varying degrees of bone spurs (osteophytes) and bony bridges across the vertebral disc space (Kranenburg et al. 2011). These lesions are not congenital, but are a degenerative change commonly related to age (Larsen and Selby 1981; Kranenburg et al. 2011) that may also form as a result of traumatic fractures or abnormal vertebral anatomy and alignment (Morgan et al. 1989). Two specimens (F75 and M152) had severe spondylosis that may have affected their spinal mobility. In sum, wolves on Isle Royale suffer from a suite of vertebral changes including types that can impair mobility and fitness, which are rarely observed in outbred wolves.

### Genome-wide patterns of variation shaped by demographic history

To assess how demographic history has shaped spatial patterns of diversity across the genome, we calculated per-site heterozygosity in non-overlapping 1 Mb windows within each individual (Fig. 2, S2). Qualitatively, we observed three distinct patterns of genome-wide heterozygosity: 1) genomes with high heterozygosity throughout (Fig. 2A, B); 2) genomes with low heterozygosity throughout (Fig. 2C, D); and 3) genomes with a sawtooth-like pattern characterized by regions of high heterozygosity interspersed by long runs of homozygosity (ROH) devoid of variation (Fig. 2E, F). We observed high heterozygosity across the entire genome in individuals derived from outbred populations with large long-term effective population sizes, such as lowland Chinese (Xinjiang) and Minnesota wolves (Gray et al. 2009; vonHoldt et al. 2011, 2016; Fan et al. 2016). Conversely, low heterozygosity across the genome was associated with a history of isolation and small long-term effective population size, such as in Ethiopian and Tibetan wolves (Gottelli et al. 1994, 2004; Zhang et al. 2014; Fan et al. 2016). Finally, the sawtooth-like pattern characterized by long ROH was observed in individuals with a history of recent inbreeding, as in the Isle Royale and Mexican wolves (Hedrick et al. 2014; Fredrickson et al. 2007). In some cases, ROH spanned entire chromosomes, consistent with the predictions of a prior simulation study based on the pedigree of the Isle Royale wolf population (Hedrick et al. 2016).

**Fig. 2.**
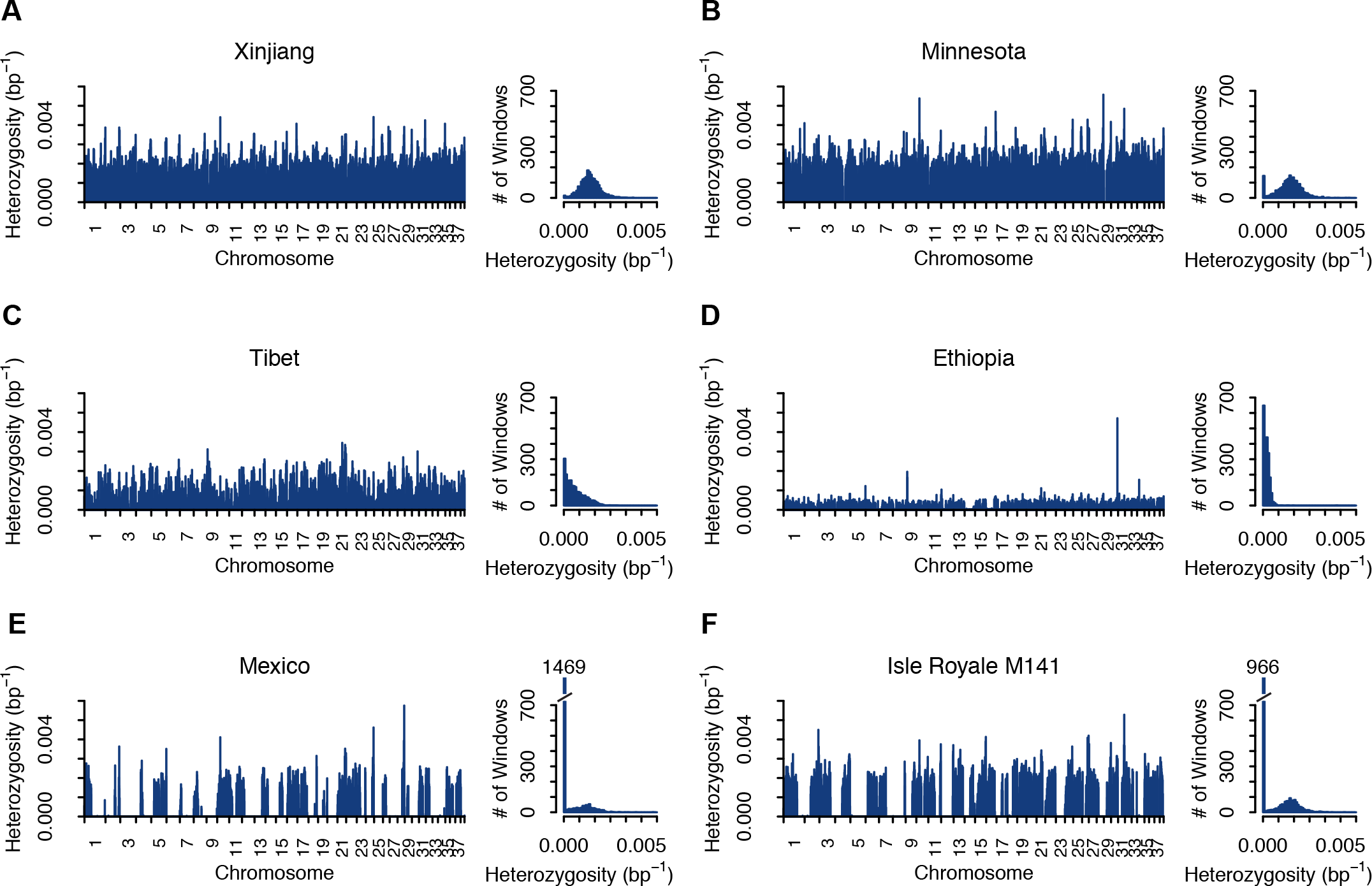
Distributions of heterozygosity across the genome. In each panel: left, example barplots showing per-site heterozygosity in non-overlapping 1 Mb windows across the autosomal genome; right, histograms of per-window heterozygosity. (**A**, **B**) The Xinjiang and Minnesota wolves represent large, outbred populations. (**C**, **D**) The Tibetan and Ethiopian wolves represent small, isolated populations. (**E**, **F**) The Mexican and Isle Royale wolves represent populations with recent inbreeding. See Fig. S2 for plots of all individuals.

Pedigree-based inbreeding coefficients (F_PED_) are predictions of the proportion of the genome that is contained within ROH (F_ROH_), which are chromosomal segments inherited identically by descent (IBD) from a common ancestor. We calculated that 23-47% of the Isle Royale wolf genomes were contained within ROH greater than 100 kb in length, which is significantly greater than F_ROH_ in Minnesota wolves, who carry 12-24% of their genomes within ROH (Mann-Whitney U (MWU) test, *p*=3.23 × 10^−4^) (Fig. 3). Our dataset includes several individuals for which we have no genealogical records (F25, F55, M61, F67), since the pedigree only includes Isle Royale wolves genotyped between 1998-2013. Among the pedigreed wolves (*n*=7), F_PED_ values ranged from 0 to 0.375. We used linear regression to evaluate the relationship between F_PED_ and F_ROH_ in the Isle Royale wolves. Pedigree-based inbreeding coefficients (F_PED_) often underestimate the true proportion of the genome contained within ROH (F_ROH_) because they fail to capture ancient or background levels of inbreeding, particularly when the pedigree is shallow, as is the case for Isle Royale wolves. The slope of the regression was shallow (0.426, *p*=0.0238) and the correlation between F_PED_ and F_ROH_ was modest (0.607, *p*=0.024), reflecting the limitations of the pedigree and the small number of samples. Nonetheless, F_PED_ (where known) always underestimated F_ROH_ (intercept=0.297, *p*=9.95 × 10^−5^) (Fig. 3A), consistent with a prior history of inbreeding not captured by the pedigree. In other words, recent severe inbreeding in Isle Royale wolves has sharply increased homozygosity across the genome, beyond the expected values suggested by the known pedigree.

**Fig. 3.**
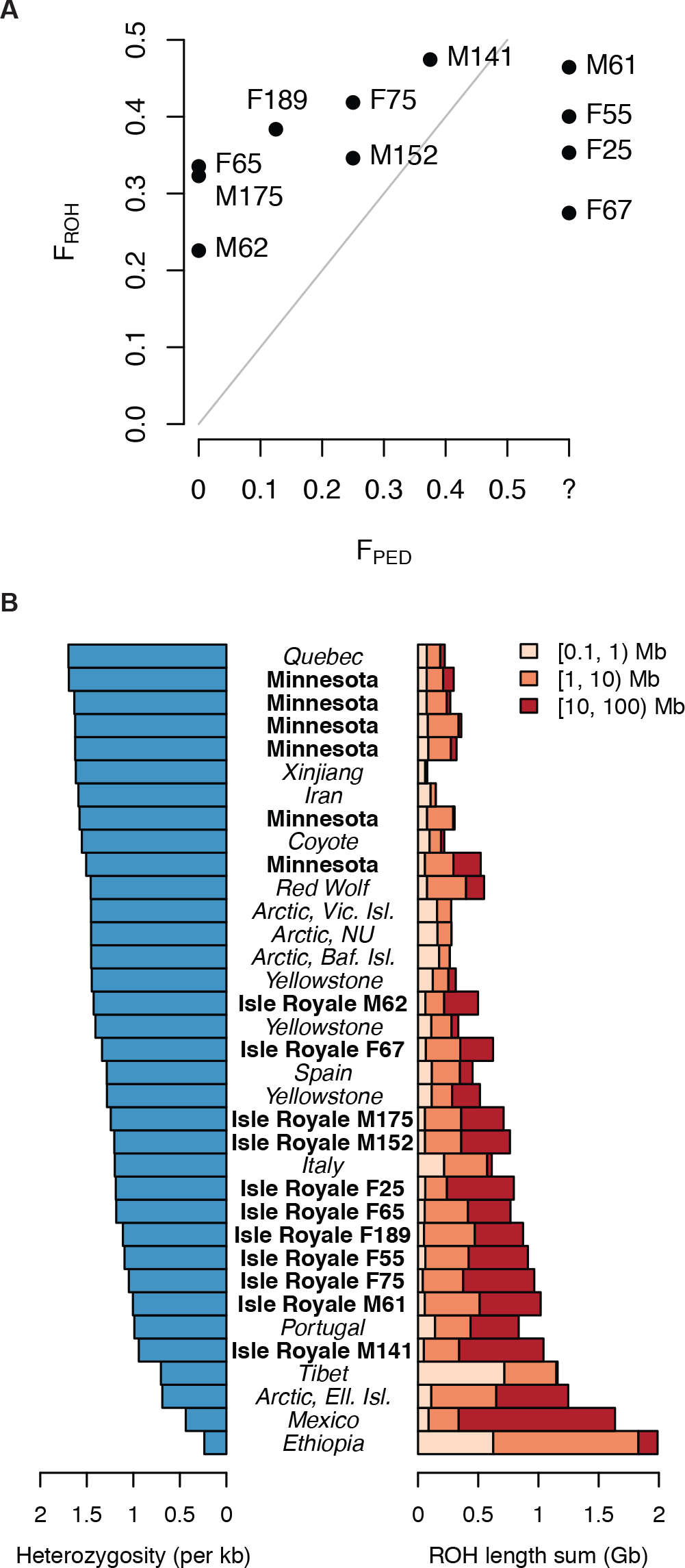
ROH and the correspondence with FPED and genome-wide heterozygosity. (**A**) F_ROH_ is the proportion of the genome contained within ROH ≥ 100kb. F_PED_ values were calculated from the pedigree of Hedrick et al. (2014) (Fig. 1A). The grey line shows the diagonal. (**B**) Left: Per-site autosomal heterozygosity across the autosomal genome. Samples are ordered by decreasing heterozygosity from top to bottom. Right: Summed lengths of short (0.1 Mb ≤ ROH < 1 Mb), medium (1 Mb ≤ ROH < 10 Mb), and long (10 Mb ≤ ROH < 100 Mb) ROH per individual.

Reduced heterozygosity in Isle Royale wolf genomes is due to the presence of extremely long ROH (>10 Mb). Overall, genome-wide heterozygosity in Isle Royale wolves (0.94-1.43 heterozygotes per kb) is 11-41% lower than the mean heterozygosity of wolves from nearby mainland Minnesota (1.60 heterozygotes per kb) (Fig. 3B). Although total heterozygosity within the genome correlates with the total amount of the genome within ROH, it does not reflect the distribution of ROH lengths, which is shaped by demographic history (Fig. 3B). The expected genetic map length of an IBD tract is inversely proportional to the number of generations (ie. the number of recombination events) since the common ancestor of the chromosomal segment (Thompson 2013). Thus, long ROH indicate recent inbreeding, whereas shorter ROH indicate ancient common ancestry. Isle Royale wolves contain 14-35 long ROH (>10 Mb) per individual, whereas Minnesota wolves contain just 1-9 long ROH per individual (Fig. S3). Long ROH are also prevalent in the genomes of the inbred Mexican wolf and the Ellesmere Island wolf, the latter suspected but not previously known to be inbred (Carmichael et al. 2008) (Fig. 3B, S3). The Mexican wolf genome derives from the highly inbred Ghost Ranch lineage, before measures were taken in the Mexican wolf captive breeding program to decrease the level of inbreeding. The increase in long ROH, but not shorter ROH, in Isle Royale wolf genomes is consistent with their recent descent from mainland wolves, which bear few ROH of any length and have high heterozygosity across their entire genomes.

In contrast, the Tibetan and Ethiopian wolf genomes contain many shorter ROH, resulting from long-term small effective population sizes, but few long ROH, suggesting no recent inbreeding (Fig. S3). Notably, these wolves have lower genome-wide heterozygosity than the Isle Royale wolves (Fig. 3B). The Tibetan wolf contained the highest number of short ROH (0.1-1 Mb) and the Ethiopian wolf genome contained the highest number of medium-sized ROH (1-10 Mb) (Fig. S3). Both the Ethiopian and Tibetan wolves exist in small isolated populations (Gottelli et al. 1994, 2004; Zhang et al. 2014; Fan et al. 2016). However, neither the Tibetan wolf nor the Ethiopian wolf is known to suffer from inbreeding depression. In fact, Ethiopian wolves are thought to practice inbreeding avoidance (Sillero-Zubiri et al. 1996; Randall et al. 2007). Since the Tibetan and Ethiopian wolves are not thought to suffer from inbreeding depression, these results imply that the intensity and timing of inbreeding are key factors modulating the risk of inbreeding depression.

### The genetic basis of inbreeding depression in Isle Royale wolves

Although the distribution of ROH number and length within a genome are informative about past inbreeding, it is still unclear how the genomic landscape of heterozygosity impacts fitness. To explore this relationship, we focused on protein-coding regions of the genome, which are more likely to directly affect fitness, and are also more amenable to functional interpretation. As the proportion of the genome contained within ROH increases, the amount of coding sequence contained within ROH increases linearly (*R*^2^=0.994, *p*<2.20 × 10^−16^) (Fig. S4). Thus, ROH are not enriched for coding regions beyond that expected for their genome-wide distribution, suggesting it is not an increase in the proportion of coding regions in ROH that causes inbreeding depression. Neither the overall homozygosity of a genome nor its coding regions is a reliable predictor of inbreeding depression, suggesting that the homozygosity of a small subset of variants have a disproportionate effect on fitness.

To assess the putative biological effects of particular mutations, we annotated variants in coding regions with respect to their impact on the encoded amino acid (eg. synonymous, non-synonymous, etc.) using the Variant Effect Predictor (VEP, McLaren et al. 2010), and further classified non-synonymous SNPs as likely to be deleterious or tolerated with the Sorting Intolerant From Tolerant (SIFT, Kumar et al. 2009) algorithm, which predicts whether missense mutations are likely to be deleterious or tolerated on the basis of amino acid conservation at a site across taxa. We then classified all variants as putatively damaging or benign. Here, synonymous and tolerated missense mutations comprise the benign group, whereas deleterious missense mutations, and mutations that disrupt splice sites or start or stop codons comprise the damaging group. Studies in humans have revealed that long ROH are enriched for homozygous deleterious variants (Szpiech et al. 2013). We did not observe enrichment of deleterious variants within ROH in our dataset (Fig. S5). However, the comparison of our results with those obtained in humans is complicated by biological differences (e.g. highly divergent demographic histories sampled within our study) and technical differences between studies (e.g. different methods for identifying and classifying ROH, potential misclassification of deleterious versus benign variants in wolves).

To control for long-term demographic history and to gauge how recent inbreeding has impacted patterns of variation in coding regions in Isle Royale wolf genomes, we focused specifically on comparing Isle Royale genomes to those of the mainland Minnesota wolves, which have a shared history until the recent founding of the Isle Royale population. Overall, both damaging and benign variants in Isle Royale wolves have shifted from the heterozygous to the homozygous state (Fig. 4A, B). The proportion of damaging homozygous genotypes was 4.07% higher (MWU *p*=3.23 × 10^−4^) in Isle Royale compared to the mainland, and 3.84% higher (MWU *p*=6.46 × 10^−4^) for tolerated homozygotes (Fig. 4B). However, the total proportion of derived alleles per genome across the Minnesota and Isle Royale wolves was unchanged for both damaging and benign mutations (MWU *p*>0.52) (Fig. 4C), consistent with population genetic theory, as inbreeding distorts genotype frequencies rather than allele frequencies. Further, because the founding of the Isle Royale population was very recent (~16 generations ago, assuming 4.2 years/generation; Peterson et al. 1998) and its numbers have remained low, an accumulation of new deleterious variants entering the population through mutation or extensive drift of existing weakly deleterious variants would not be expected. In other words, inbreeding depression in Isle Royale wolves cannot be explained by an increase in the number of derived deleterious variants per genome, which may be proportional to the additive genetic load. Rather, it must be accounted for by the increased homozygosity of deleterious variants.

**Fig. 4.**
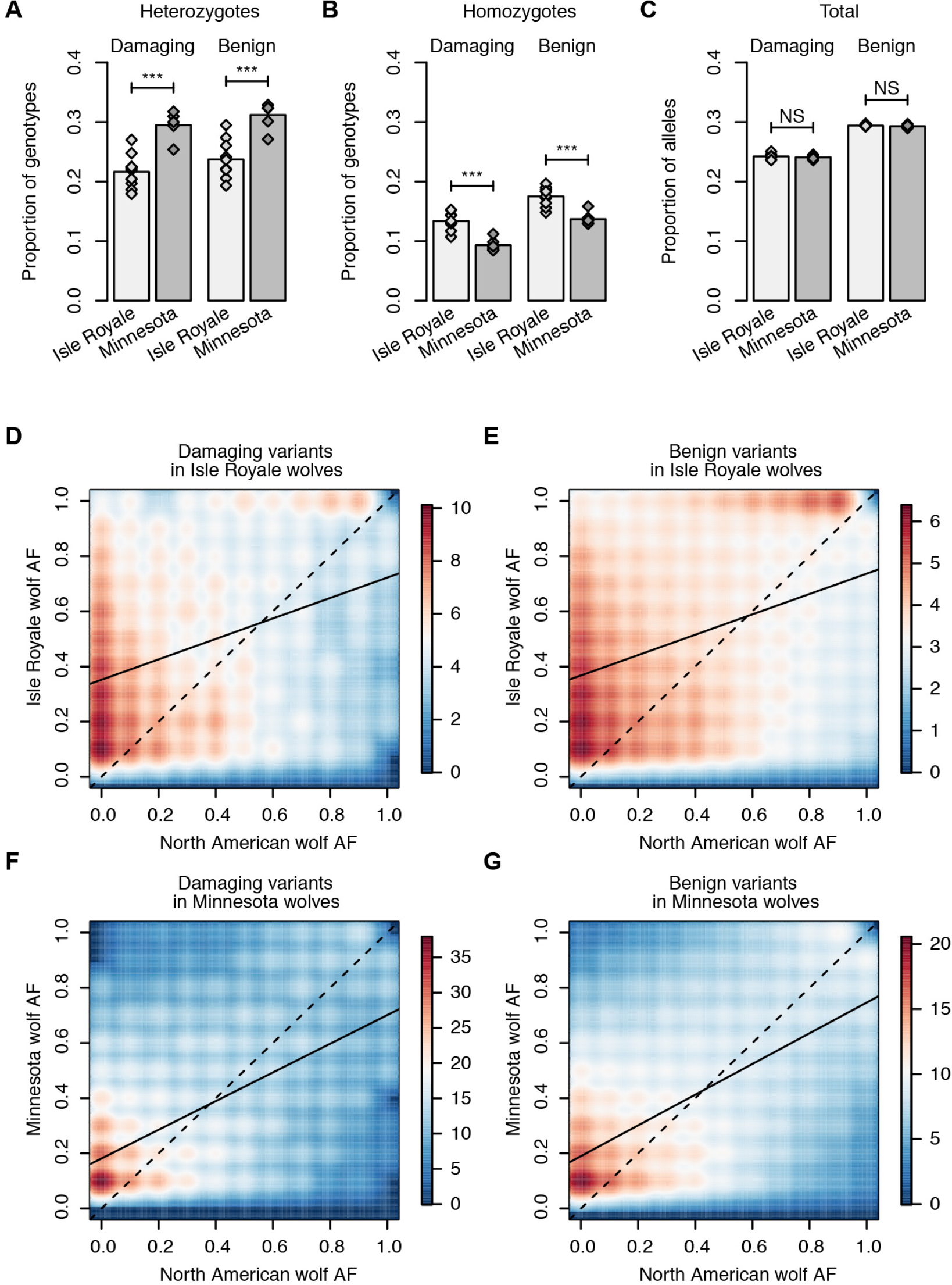
Genotype and allele frequencies in Isle Royale versus mainland Minnesota wolves. (**A**) Inbred Isle Royale wolves contained significantly fewer heterozygotes and, (**B**) significantly more homozygotes than Minnesota wolves, for both damaging and benign SNPs. (**C**) The total number of derived alleles is unaffected by recent inbreeding. Significance codes: ***, *p* < 0.001; NS, not significant. (**D-G**) Two-dimensional allele frequency spectra showing the correlation in derived allele frequencies (AF) between outbred North American wolves (Quebec, Yellowstone, Canadian Arctic excluding Ellesmere Island) and Isle Royale or Minnesota wolves, for variants present in Isle Royale or Minnesota wolves. All sites were down-sampled to include exactly five individuals from each group. Color represents the density of points (see legends). The dashed line represents the diagonal, whereas the solid line represents the linear regression line (see Table S3). (**D**) 3,697 sites, (**E**) 48,326 sites, (**F**) 5,216 sites, (**G**) 63,893 sites.

Even strongly deleterious recessive alleles are expected to be present in standing genetic variation, segregating at low frequencies in large populations where drift is minimal and inbreeding is rare. In contrast, variants that are at high frequency in large populations of outbred wolves are not likely to be strongly deleterious. Thus, in the absence of appreciable gene flow with the mainland, we predicted that strongly deleterious recessive alleles carried in the founder genomes could have attained high frequency within the Isle Royale population. This phenomenon has been observed in the increased prevalence of rare genetic disorders in founder populations of humans (reviewed by Sheffield et al. 1998) and purebred dogs (reviewed by Sutter and Ostrander 2004). We compared the frequencies of segregating variants in Isle Royale wolves and mainland Minnesota wolves to those of outbred North American wolves by constructing two-dimensional allele frequency spectra, and performing linear regression to assess correlations between populations (Fig. 4D-G, Table S3). For this analysis, all groups were down-sampled to 10 chromosomes (5 individuals) each. We found that variants with low frequency in outbred North American wolves are also typically at low frequency in mainland Minnesota wolves, consistent with weak drift and efficient selection (Fig. 4F, G, Table S3). In contrast, non-synonymous variants in Isle Royale wolves had higher frequencies due to the effects of isolation and high relatedness among individuals (Fig. 4D, E). Further, for both damaging and benign variant classes, derived allele frequencies in Isle Royale had lower correlations with allele frequencies in outbred wolves compared to Minnesota wolves, consistent with our prediction that the founder effect and inbreeding in Isle Royale wolves allowed damaging variants to attain high frequency (*p*<2.22 × 10^−16^).

We predicted that these damaging variants might account for the high incidence of vertebral anomalies observed in Isle Royale wolves. We used the following criteria to identify candidate deleterious variants underlying the phenotypes of Isle Royale wolves in our dataset: 1) homozygous and located within ROH in the affected individuals (F65, F75, M152, M175, F189) but heterozygous or absent in the unaffected individual (M61); and 2) low frequency (<10%) among other gray wolves (*n*=21). 263 genes containing such mutations were found in at least one affected individual, and two of these genes contained mutations in all five affected individuals: *RTTN* and *SCUBE2*. Only *RTTN* (rotatin) is known to be associated with abnormal phenotypes, and notably affects vertebral development. *RTTN* is a large, highly conserved gene with 49 exons, spanning 144 kb on chromosome 1 of the dog genome.

Using a mouse model, researchers have found that *RTTN* plays an essential role in early embryonic development, specifically in left-right specification, embryo turning, and notochord formation (Faisst et al. 2002). Embryos with *RTTN* knocked out were inviable, but developed normally with one functional gene copy. The five Isle Royale wolves with vertebral malformations are homozygous for a C to T transition that converts a leucine residue to phenylalanine in exon 11 of *RTTN*. The SIFT score for this mutation is 0, indicating very strong conservation at this site, and that this mutation is therefore predicted to have a deleterious effect. No other homozygotes for this mutation were present in our dataset, whereas three of the six other Isle Royale wolves and two of the Minnesota wolves were heterozygotes. Of the 263 candidate genes we identified in the affected individuals, 10 are associated with the Human Phenotype Ontology (HPO) term “abnormality of the vertebral column” (HP:0000925), but were not shared across all five affected individuals. The 10 genes are *ABCC6*, *CAPN1*, *ELN*, *ERCC1*, *MLXIPL*, *PSAT1*, *TCTN2*, *TERT*, *ENSCAFG00000001588* (ortholog of *SLC52A2*), and *ENSCAFG00000006532* (ortholog of *DCHS1*) (Fig. S6). Morphogenesis is a complex process, and the variation in phenotypes within the Isle Royale wolves suggests the involvement of multiple genes.

### Testing models for the mechanistic basis of inbreeding depression

We hypothesized that the reason inbreeding depression afflicts Isle Royale wolves, but not other populations with a long-term history of small population size and isolation, may be due to differences in the prevalence of severely deleterious recessive alleles in the mainland source population combined with recent inbreeding in Isle Royale. To test this hypothesis, we conducted simulations in SLiM (Haller and Messer 2016) under a two-population model incorporating estimates of the long-term effective population sizes of outbred North American wolves (N_e_=17,350) and Tibetan wolves (N_e_=2,500), and the estimated divergence time between Old and New World wolves of 12,500 years (Fig. 5A) (Fan et al. 2016). We simulated diploid individuals containing genomes of 1,000 “genes” that accumulated neutral and deleterious mutations, in order to compare the number of mutations per genome in each population. Deleterious mutations were categorized as weakly (0<N_e_*s*≤10), moderately (10<N_e_*s*≤100), and strongly (N_e_*s*>100) deleterious, where N_e_ corresponds to the size of the ancestral population before the North American and Tibetan populations diverged (N_e_=45,000). We conducted one set of simulations in which all mutations were additive (*h*=0.5), and one in which all were recessive (*h*=0), to explore the effects of dominance. The MWU test was used to evaluate statistical significance in comparisons between the two populations. Here, the North American wolf population represents the source population for Isle Royale wolves, which only became isolated fewer than one hundred years ago, whereas the Tibetan population represents a population with long-term small effective population size. We predicted that the larger North American population would contain more strongly deleterious recessive mutations per individual relative to the Tibetan population. These mutations in particular would severely compromise fitness in an individual that inherits two copies from a common ancestor through inbreeding.

**Fig. 5.**
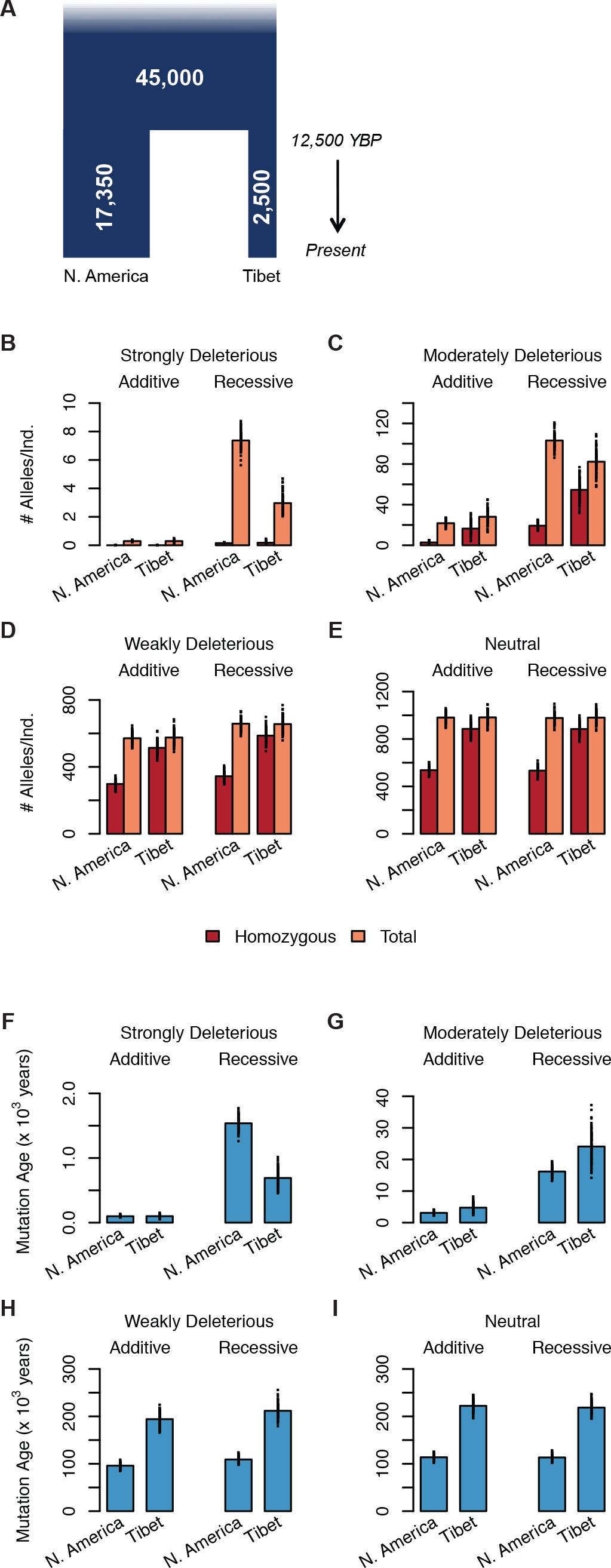
Model and results from simulations of deleterious variation. (**A**) Demographic model used to simulate the expected number and age of mutations in a large population (N. America, 17,350 individuals) versus a small population (Tibet, 2,500 individuals). Both populations split from a large ancestral population (45,000 individuals) 12,500 years before present (3-year generation time). Population sizes and split time from Fan et al. (2016). Model not drawn to scale. (**B-I**) Results from simulations, grouped according to dominance and selection coefficients. Additive: *h*=0.5, recessive *h*=0; strongly deleterious, N_e_*s*>100; moderately deleterious, 100≥N_e_*s*>10; weakly deleterious, 10≥N_e_*s*>0; neutral: N_e_*s*=0. (**B-E**) The average number of homozygous and total alleles per individual. (**F-I**) The average ages of segregating mutations in each population.

Our simulations confirmed that the total number and the homozygosity of deleterious alleles per individual vary between the larger North American and the smaller Tibetan populations. Importantly, different patterns were observed depending upon whether mutations were additive or recessive. In simulations with additive mutations, the overall number of deleterious alleles per individual was slightly higher (+1.87%) in the smaller Tibetan population relative to the larger North American population (*p*=1.84 × 10^−2^). This difference is due to the accumulation of moderately deleterious alleles in the Tibetan population (+34%, *p*=2.49 × 10^−11^), as there was no significant difference in the numbers of strongly deleterious or weakly deleterious additive alleles between the two populations. Thus, in an additive model, selection against strongly deleterious alleles is not hindered by drift (Fig. 5B), but moderately deleterious alleles accumulate under stronger drift (Fig. 5C), leading to a slight increase in the overall number of deleterious alleles in smaller populations.

However, in simulations with recessive mutations, a very different pattern emerged. Here, the overall number of deleterious alleles per individual was higher in the larger North American population (+3.58%, *P*=1.42 × 10^−6^). Although the per-individual number of weakly deleterious alleles, which make up the vast majority of deleterious mutations (>90%), was approximately equal in the two populations (Fig. 5D), individuals from the larger North American population had sharply elevated numbers of strongly (+59.2%, *P*=2.56 × 10^−34^) and moderately (+23.4%, *P*=3.04 × 10^−24^) deleterious recessive alleles (Fig. 5B, C) relative to the smaller Tibetan population. Furthermore, the mean age of segregating strongly deleterious recessive mutations was 2.22-fold higher in the North American population, compared to a mean age of 1,199 years in the Tibetan population (*P*=2.56 × 10^−34^) (Fig. 5F), indicating that these mutations persist over longer time periods while they remain hidden from selection as heterozygotes in the larger population. In contrast, segregating mutations in all other categories tended to be younger in the North American population (Fig. 5G-I), as a consequence of fewer new mutations entering the Tibetan population each generation due to its smaller size.

In sum, we found that a smaller population, such as Tibetan wolves, has fewer strongly deleterious recessive alleles, but that these mutations persist in large populations, such as North American wolves. Inbreeding would therefore produce more individuals homozygous for strongly deleterious mutations in individuals drawn from a historically large population compared to individuals drawn from a historically smaller population. Because Isle Royale wolves were recently founded from a large mainland population of wolves in the Great Lakes region, and then experienced extensive inbreeding, our simulations suggest they should have more homozygous strongly deleterious recessive mutations, resulting in increased inbreeding depression.

## Discussion

The persistence of wolves on Isle Royale was once used to support the claim that a very small population in isolation may persist, and even thrive, without succumbing to genetic deterioration (Mech and Cronin 2010). During the past few decades, however, Isle Royale wolves have experienced a precipitous decline following generations of inbreeding and physical degeneration (Räikkönen et al. 2009; Hedrick et al. 2014). Although inbreeding depression was not the sole determinant, it has undoubtedly played a role in the collapse of the population, along with stochastic demographic and environmental events, such as periodic disease outbreaks, severe winters, and the drowning of three wolves in a flooded abandoned mine shaft in 2011 (Hedrick et al. 2014). The genomes of Isle Royale wolves bear the hallmarks of their extreme demographic history, characterized by extensive ROH, in some cases spanning whole chromosomes, leading to a marked increase in the homozygosity of deleterious variants.

Notably, wolves from other populations with more homozygous genomes, but with shorter ROH, are not known to be afflicted by inbreeding depression. The absence of inbreeding in Tibetan and Ethiopian wolves is not definitive, however, and future research is needed to determine whether low genetic diversity and high homozygosity of deleterious variants in these populations is associated with abnormal phenotypes. Nonetheless, our simulations affirm that purging of strongly deleterious recessive alleles may occur in populations of moderate size and low heterozygosity, despite increases in the overall burden of deleterious variants due to the accumulation of weakly deleterious alleles. Currently, calculating the burden of strongly deleterious recessive alleles within a genome is challenging, particularly in non-model species. In humans, it has been estimated that each diploid genome carries ~1-2 recessive lethal mutations, but the number of recessive sub-lethal mutations that compromise fitness is likely to be much higher (Gao et al. 2015). Thus, the risk of inbreeding depression is higher for genomes recently originating from a historically large population, such as Isle Royale wolves, as they carry a greater burden of strongly deleterious recessive mutations. These strongly deleterious recessive mutations are carried as heterozygotes within the founders, but quickly become homozygous in the island population through inbreeding, resulting in inbreeding depression. Similar phenomena have been noted in maize, which were domesticated from historically large populations, presumably carrying many recessive deleterious alleles in the heterozygous state. Here the initial inbred lines exhibited severe inbreeding depression and reduced yield, creating the need for hybrid lines (Troyer 2006).

Our analysis of Isle Royale wolf genomes contrasts with an increasing number of studies showing an elevated burden of deleterious variants along with reduced genetic diversity in historically small or bottlenecked populations (eg. Lohmueller et al. 2008; Renaut and Rieseberg 2015; Marsden et al. 2016). We found no difference in the number of derived deleterious alleles in Isle Royale compared to the mainland Minnesota population, suggesting that the additive genetic load may be the same in both populations. Instead, the reduction in fitness on Isle Royale is due to the increased homozygosity of recessive mutations. Alternatively, the additive genetic load may be higher in the Isle Royale population, but we are unable to detect this increased load due to the difficulties of determining which amino acid changing variants are deleterious. In either scenario, our findings suggest that populations with a similar number of derived deleterious alleles and heterozygosity may still differ in their genetic load, and additional metrics should be used to quantify the load in populations.

Increased homozygosity due to severe inbreeding in Isle Royale wolves has resulted in significant morphologic defects, especially malformed vertebrae that are associated with adverse clinical symptoms in dogs. Other abnormalities have also been observed in Isle Royale wolves, including syndactyly, probable cataracts, an unusual “rope tail”, and anomalous fur phenotypes (Räikkönen et al. 2009, Peterson and Vucetich 2015). Hedrick et al. (2014) found that highly inbred wolves had low survival and reproduction relative to less inbred wolves in Isle Royale. The individual with the highest pedigree-based inbreeding coefficient and the highest homozygosity among our sequenced Isle Royale wolves, M141 (F_PED_=0.375, F_ROH_=0.47), lived only two years and did not reproduce (Hedrick et al. 2014). Another wolf, F75 (F_PED_=0.25, F_ROH_=0.42), died at four years of age while giving birth to a litter of pups presumably sired by her own father (Hedrick et al. 2014). The pedigree-based inbreeding coefficient of this litter, in which all pups showed vertebral changes and all but one possessed extra ribs, was 0.375 (Hedrick et al. 2014). Reproduction within the population ceased after 2014, and it has fallen from 30 individuals to only two over the past twelve years (Peterson and Vucetich 2017). The population of moose on Isle Royale, the main prey of Isle Royale wolves, has swelled from 450 to 1600 individuals over the same period (Peterson and Vucetich 2017). Thus the demise of wolves on Isle Royale cannot be attributed to lack of available prey.

The collapse of the Isle Royale wolf population occurred despite a reported genetic rescue and evidence of earlier sporadic migration events from the mainland. Previous genetic analysis revealed that undetected migration from the mainland may have occurred in years when the winter was cold enough for an ice bridge to form (Adams et al. 2011; Hedrick et al. 2014). However, warmer winters over the past several decades have resulted in a dramatic reduction in the formation of ice bridges, a trend that is likely to continue in a warming climate (Hedrick et al. 2014). The reported genetic rescue of the Isle Royale wolf population by a single migrant from the mainland in 1997 also appears to have been short-lived (Hedrick et al 2014). This male wolf was such a successful breeder that his genome effectively swamped the population, leading to intense inbreeding in his descendants within two generations (Fig. 1A). A similar episode occurred in the inbred Scandinavian wolf population following the arrival of an immigrant wolf in 1991, but the population has entered a period of growth following subsequent additional immigration events (Åkesson et al. 2016). A higher rate of gene flow between the mainland and the island after the Isle Royale population was established may have mitigated or postponed inbreeding depression, but the effective rate of naturally-occurring gene flow was clearly insufficient. Sustained human-assisted gene flow may therefore be the only option for the persistence of wolves on Isle Royale.

A potential alternative strategy to reduce the risk of inbreeding depression following reintroduction would be to select founders from a historically small population, where purging of strongly deleterious alleles may have already occurred. Such individuals may be effectively pre-adapted to withstand small population size, bottlenecks, and inbreeding. This strategy must be considered carefully, however, since its success requires the absence of gene flow with large populations nearby that would introduce strongly deleterious recessive alleles back into the smaller population. Given the intermittent migration that occurs with the mainland, selecting founder individuals from historically small populations would not be a viable strategy for Isle Royale wolf reintroduction. Nonetheless, the idea that founders from historically isolated populations should be selected in order to mitigate the risk of inbreeding depression is novel, but may be an approach to enhance long-term persistence of the inevitably small and isolated populations of many species in the future.

Finally, life history traits must be considered when determining the best course of action to reduce the risk of inbreeding. For example, wolves, including those on Isle Royale, typically avoid mating with close relatives through the exchange of individuals between different packs, which are usually familial units that consist of a breeding pair and its offspring (Geffen et al. 2011). Previously, Isle Royale sustained 3-4 wolf packs, hence the estimate that the long-term effective population size of wolves on Isle Royale was a mere 3.8 individuals (Peterson et al. 1998). In simulations under models incorporating ecological and demographic stochasticity, the mean time to extinction for social organisms, specifically wolves, is strongly tied to the number of social groups rather than the number of individuals (Vucetich et al. 1997). Thus, a minimum number of wolves to sustain multiple packs, and therefore maximize the number of breeding individuals is essential for population persistence.

A vigorous ongoing discussion concerns the restoration of a healthy wolf population on Isle Royale by introducing wolves from the mainland (Vucetich et al. 2016). In the absence of recurring immigration, whether managed or not, the fate of a restored population is grim, given the inevitability of inbreeding on Isle Royale and its proven detrimental outcome. On the other hand, wolf predation is an important top-down influence on Isle Royale, and its absence threatens the stability of the island ecosystem (Peterson et al. 2014). The collapse of the Isle Royale wolf population demonstrates the critical importance of maintaining effective population sizes large enough to allow selection to remove strongly deleterious variants. Although it is too late to resurrect a population from the lone remaining pair of wolves on Isle Royale, we show that the demise of this iconic population provides new lessons for a potential reintroduction, as well as guidance for the management of other species or populations to minimize the risk of inbreeding depression.

## Materials and Methods

### Samples and sequencing

DNA from Isle Royale wolves was extracted from blood samples archived at Michigan Technological University. DNA samples with high quality and high molecular weight were selected for sequencing. DNA from Minnesota and Canadian Arctic wolves was extracted from blood and tissue samples from the archive of Dr. Robert Wayne that were used in previous studies (vonHoldt et al. 2011, Schweizer et al. 2016). Whole genome sequencing was performed on an Illumina HiSeq4000 at the Vincent J. Coates Genomics Sequencing Laboratory at UC Berkeley. Previously sequenced genomes with high coverage were downloaded from the NCBI Short Read Archive (see Table S1).

### Read processing and alignment

A pipeline adapted from the Genome Analysis Toolkit (GATK, McKenna et al. 2010) Best Practices Guide was used to process raw reads prior to genotype calling. Briefly, paired end raw sequence reads, 150 bases in length, were aligned to the domestic dog reference genome, canFam3.1, using bwa MEM (Li 2013), before removal of PCR duplicates and low quality reads. Lacking a database of known variants, the bootstrapping method of base quality score recalibration as recommended by GATK was performed by calling raw genotypes with GATK UnifiedGenotyper (minimum base quality Phred score 20), and using these variants as input for recalibration with BaseRecalibrator. This process was repeated three times to reach convergence between reported and empirical quality scores.

### Genotype calling and filtering

Joint genotype calling was performed with GATK HaplotypeCaller. Genotypes were filtered for quality and depth, leaving only high quality biallelic single nucleotide polymorphisms. Only genotypes with at least six supporting reads and high quality (minimum Phred score of 20) were included. An excess depth filter, set at the 99^th^ percentile of depth for each sample, was also used. Variant sites were then filtered on the following criteria: sites failing the recommended GATK hard filters were excluded, as well as sites with excess depth (>99^th^ percentile for total depth across all samples), low Phred score (QUAL<30), more than 20% missing data, excess heterozygosity (>50% of individuals heterozygous), or sites found within repeat regions and CpG islands (coordinates from Marsden et al. 2016).

### Variant annotation

Variant sites were annotated with the Ensembl VEP (version 87) with SIFT enabled (Kumar et al. 2009; McLaren et al. 2010). SIFT determines whether a nonsynonymous mutation is likely to be damaging or benign on the basis of phylogenetic constraint on an amino acid within a protein alignment. We grouped variants in protein-coding regions into “damaging” and “benign” classes. Damaging variants included nonsynonymous variants classified as “deleterious” by SIFT (score <0.05) and variants that disrupted splice sites, start codons, or stop codons. Benign variants included nonsynonymous variants classified as “tolerated” by SIFT (score ≥0.05) and synonymous mutations. Alleles were polarized as derived or ancestral with respect to the gray fox and African golden wolf genomes.

### Phylogenetic analysis and cladogram construction

A cladogram representing the relationships between 20 genomes in this study was constructed using SNPhylo (Lee et al. 2014). Where multiple individuals were available from a single population, one individual was chosen at random for inclusion in the tree (Minnesota: RKW119, Isle Royale: CL141, Yellowstone: 569F). SNPs were pruned for linkage disequilibrium (threshold of 0.2), and SNPs with minor allele frequency <0.1 or with missingness above 10% were excluded. The tree was constructed with the 28,651 remaining SNPs. 1,000 bootstrap replicates were performed.

### Morphological analysis

Skeletons from six of the eleven Isle Royale wolves included in this study were retrieved from storage at Michigan Technological University, photographed, and assessed for morphological anomalies as described in Räikkönen et al. 2006, 2009. Skeletons were obtained from carcasses collected post mortem. The vertebral column and rib variation was evaluated. In some cases, specimens were missing a few vertebrae or ribs; the list of examined wolves is noted in Table 1. A litter of eight newborn pups from F75 was also examined through radiographic analysis.

### Calculation of genome-wide heterozygosity

In this study, we calculated heterozygosity as the number of heterozygous genotypes divided by the total number of called genotypes within a single individual. For each individual, we calculated heterozygosity for the entire autosomal genome, as well as in non-overlapping 1-Mb windows across the autosomes. Windows where more than 80% of sites failing filters or missing were excluded.

### Identification and analysis of ROH

ROH were identified using VCFtools (Danecek et al. 2011). ROH spanning regions with fewer than 50 variant sites were excluded. The amount of protein-coding sequence within ROH was determined by calculating the overlap between the coordinates of ROH within each individual and the coordinates of protein-coding exons download from Ensembl Biomart (version 87). To test for enrichment of homozygous deleterious variants within ROH, we followed the method of Szpiech et al. 2013. The fraction of the genome within ROH and the fraction of damaging and benign homozygous genotypes inside ROH was calculated in each individual. Following Equation 10 of Szpiech et al. 2013, we fit linear models to test whether variant impact (benign versus deleterious) and F_ROH_ were significant predictors (β_2_ and β_3_, respectively) for the proportion of nonreference homozygotes within ROH.

### Identification of candidate genes underlying Isle Royale phenotypes

Candidate mutations underlying the abnormal phenotypes observed within our sample of Isle Royale wolves satisfying the following criteria were identified. Mutations had to be classified as damaging (missense mutations classified as deleterious by SIFT as well as mutations disrupting splice sites or start/stop codons) and passing all quality control filters described above. Mutations had to be homozygous and contained within ROH of 100 kb or more in the affected wolves (F65, F75, M152, M175, F189), but not homozygous for the derived allele in the unaffected wolf (M61). Finally, only mutations with low frequency (<10%) in 21 other gray wolves (wolves from Minnesota (6), Canadian arctic (4), Yellowstone (3), Quebec, Mexico, Portugal, Spain, Italy, Iran, Tibet, Xinjiang) were considered. Alleles were polarized as ancestral or derived with respect to the gray fox and African wolf outgroups. Genes containing mutations satisfying all criteria were extracted and associated with HPO terms (2017-10-05 release) using gProfileR (r1741_e90_eg37 release, Reimand et al. 2011).

### Simulations of neutral and deleterious variation

Simulations were carried out in SLiM (version 2.4.2, Haller and Messer 2016) under a divergence model with parameters estimated by Fan et al. (2016). Each simulated individual consisted of a diploid 1 Mb genome, with a simple architecture of 1,000 “genes” carried on 38 chromosomes proportional to chromosome lengths in the dog genome. Each gene consisted of a contiguous 1 kb sequence that accumulated mutations at a rate of 1 × 10^−8^ per site per generation. Selection coefficients for deleterious mutations were drawn from the distribution of fitness effects inferred from a large sample of humans by Kim et al. (2017). 70% of mutations were deleterious, and the remaining 30% were neutral (*s*=0). Each simulation began with a burn-in period of 450,000 (10 × N_e_) generations to allow the ancestral population to reach equilibrium. Recombination was permitted at single base positions between each gene at a rate of 1 × 10^−3^ per site per generation, to simulate the effective rate of crossing over that would occur in 100 kb noncoding regions between each gene. At the end of each simulation, the average number of alleles per individual and the average age of segregating mutations were calculated for weakly (0<N_e_*s*≤10), moderately (10<N_e_*s*≤100), strongly (N_e_*s*>100) deleterious, and neutral mutations (N_e_*s*=0). We performed 100 replicates in which mutations were additive (*h*=0.5) and 100 replicates in which mutations were completely recessive (*h*=0.0), to examine the effects of dominance.

## Acknowledgements

We thank Philip W. Hedrick for helpful discussions in preparation of this manuscript. This work used the Vincent J. Coates Genomics Sequencing Laboratory at UC Berkeley, supported by NIH S10 OD018174 Instrumentation Grant. This work was funded by National Institute of Health grant R35GM119856 to KEL, U.S. National Science Foundation DEB-1453041 to JAV, Isle Royale National Park (CESU Task Agreement No. P16AC00004, under Master Cooperative Agreement Number P12AC31164), the Robbins Chair in Sustainable Management of the Environment to ROP at Michigan Technological University, McIntyre-Stennis Grant USDA-Nifa-1014575.

## Author Contributions

The study was conceived of and designed by JAR, KEL, RKW, and ROP. Sample acquisition was performed by LMV, JAV, and ROP. Morphological analyses were carried out by JR. Genomic analyses and simulations were carried out by JAR. Manuscript was written by JAR. All authors contributed to manuscript revision and approved the final version. RKW and KEL jointly supervised this work.

## Data and Materials Availability

Newly generated genome sequence data were deposited in the NCBI Short Read Archive under BioProject PRJNAXXXXXX. The authors acknowledge Nathan C. Nelson from Michigan State University for providing radiographs of the wolf pups.

## Supplemental Figures and Tables

**Fig. S1.**
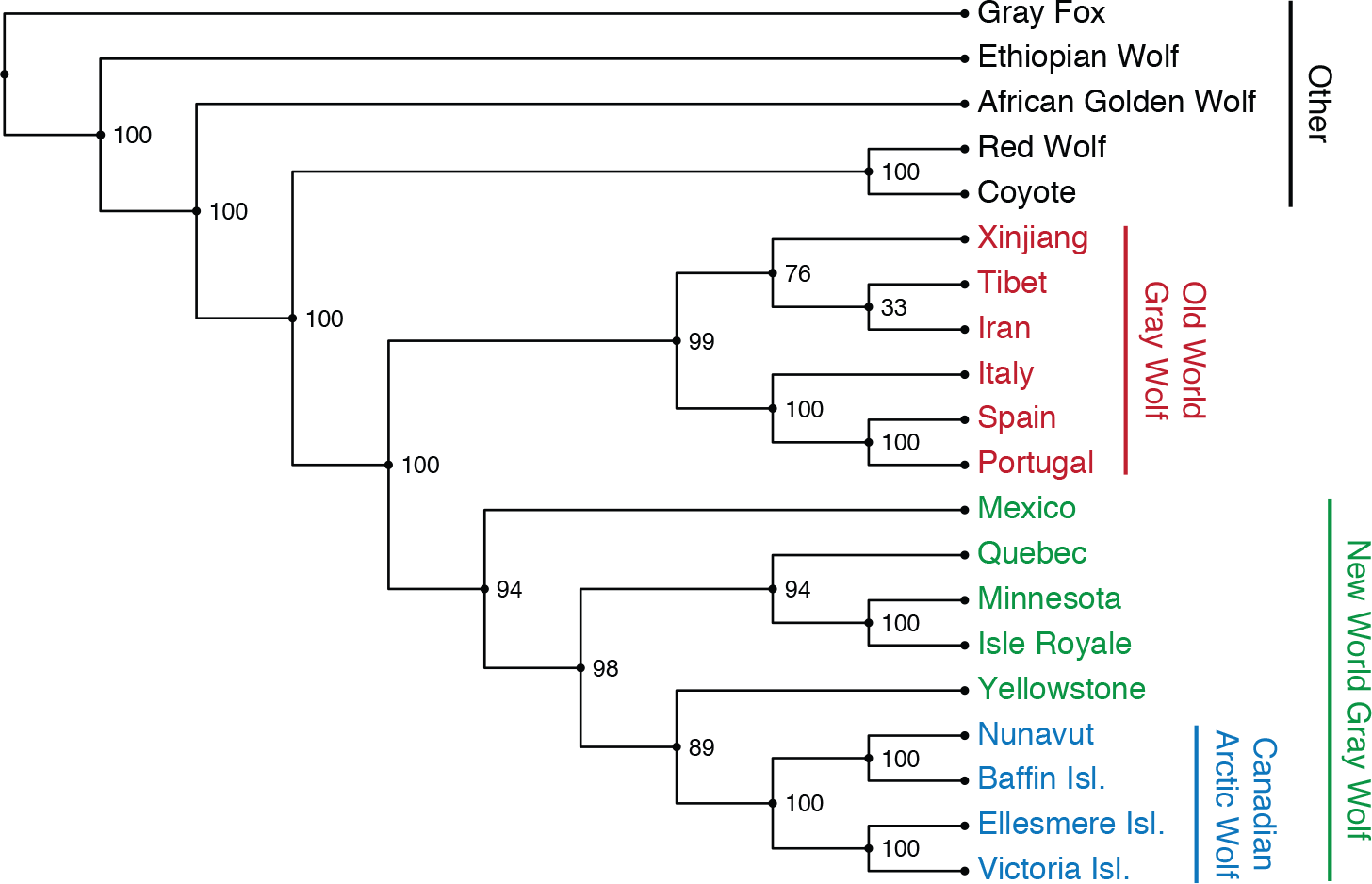
Cladogram of genome sequences in this study. A phylogeny of 20 genome sequences based on 28,651 SNPs pruned for linkage disequilibrium shows the relationships among wolf populations and sister taxa. Where multiple individuals were available from a single population, one individual was chosen at random for inclusion in the tree (Minnesota: RKW119, Isle Royale: CL141, Yellowstone: 569F). Percentage of support from 1,000 bootstrap replicates is indicated at each node.

**Fig. S2.**
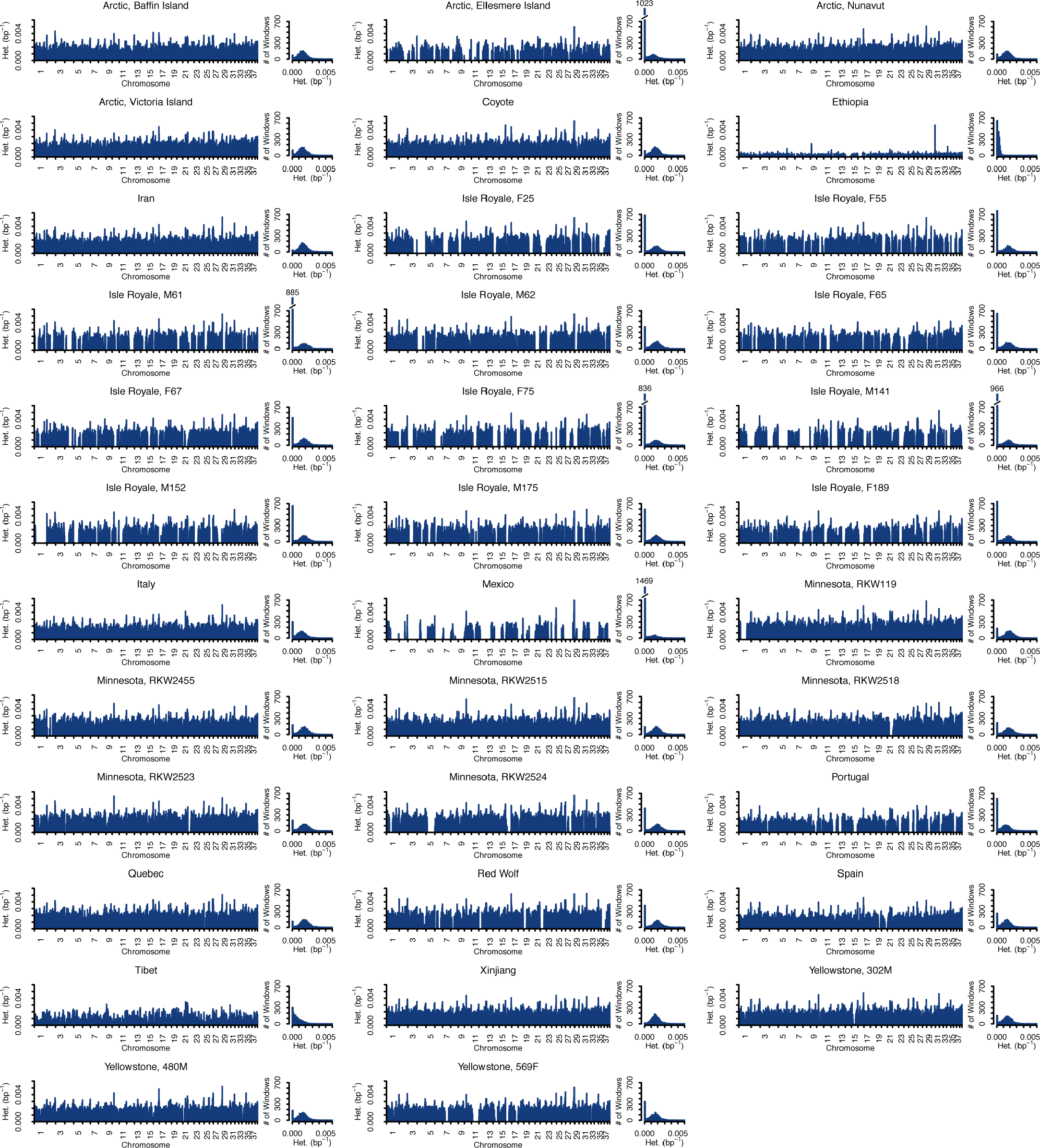
Distributions of heterozygosity in all individuals. In each panel: left, example barplots showing per-site heterozygosity in non-overlapping 1 Mb windows across the autosomal genome; right, histograms of per-window heterozygosity.

**Fig. S3.**
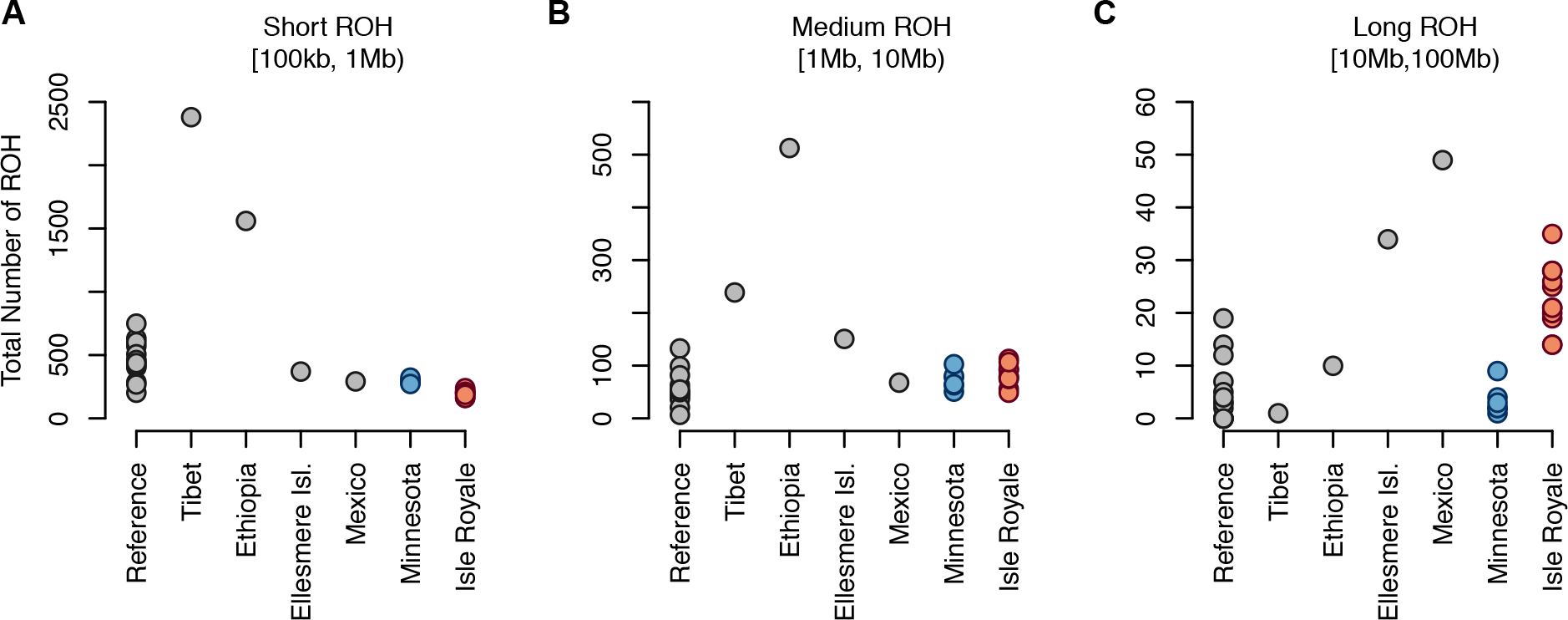
Number of ROH per individual for different ROH length categories. The number of ROH in various size classes is indicative of demographic history. “Reference” individuals include gray wolves from Canadian Arctic (except Ellesmere Island), Quebec, Yellowstone, Iran, Italy, Spain, Portugal, and Xinjiang, plus the red wolf and California coyote. (**A**) Short ROH indicate ancient inbreeding, as in the Tibetan wolf. (**B**) Medium ROH indicate ancient and historic inbreeding, as in the Ethiopian wolf. (**C**) Long ROH indicate recent inbreeding, as in the Mexican, Isle Royale, and Ellesmere Island wolves.

**Fig. S4.**
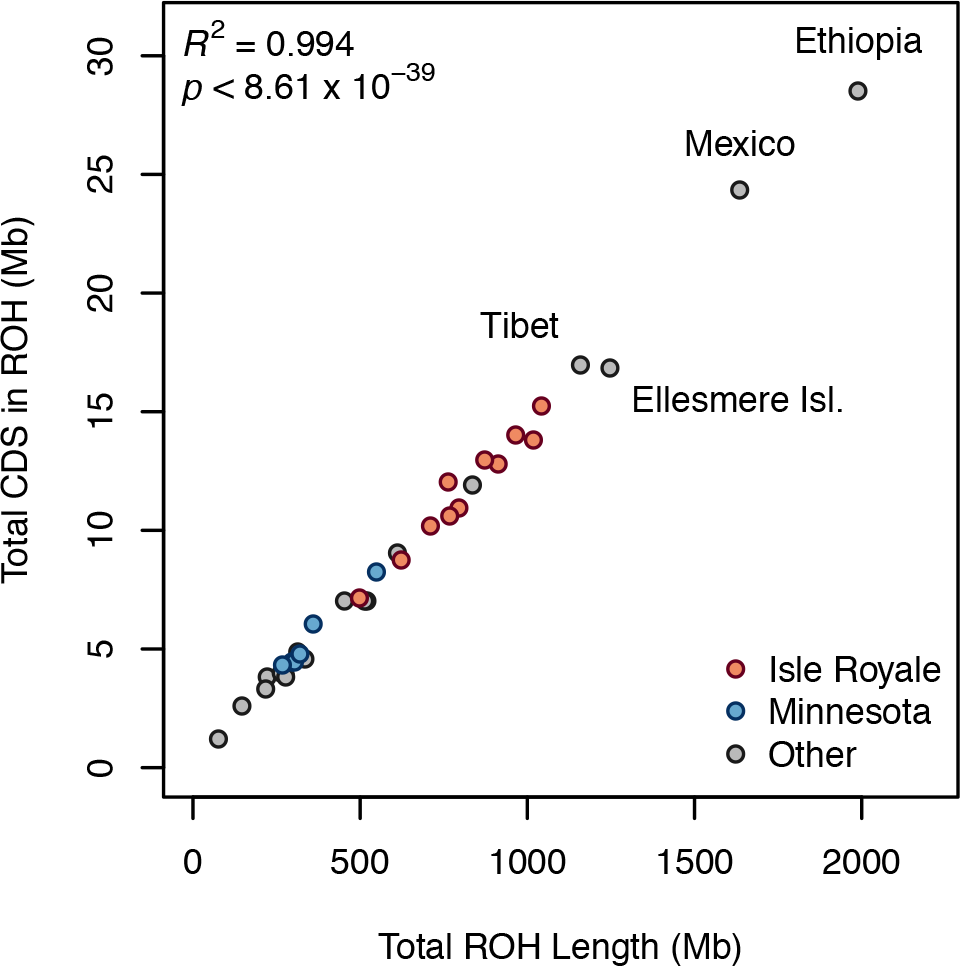
Amount of coding DNA sequence in ROH as a function of the amount of the genome within ROH. *R*^2^ correlation and *p*-value coefficients were obtained by linear regression.

**Fig. S5.**
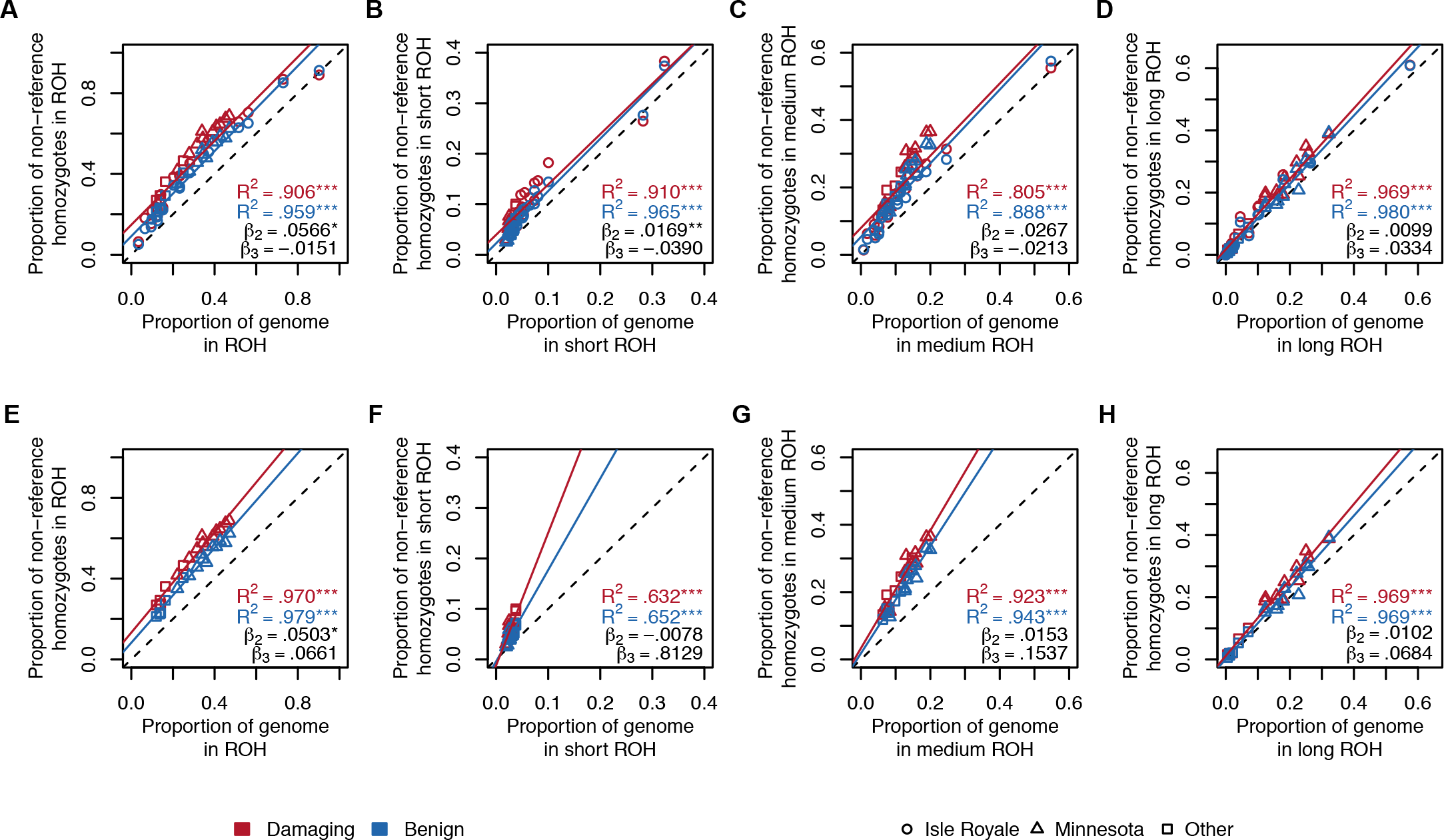
Proportion of non-reference homozygotes in ROH as a function of the proportion of the genome within ROH. Following Szpiech et al. 2013, the proportion of non-reference homozygotes was calculated for benign and damaging variants and plotted against the proportion of the genome within ROH in all 35 individuals (**A-D**) and just Isle Royale and Minnesota wolves (**E-H**). Linear regression correlation coefficients (R^2^) and their significance are indicated. The β-coefficients were calculated as in Equation 10 of Szpiech et al. 2013. β_2_ and β_3_ indicate the change in intercept and slope, respectively, for damaging variants relative to benign variants. Significance codes: ***, *p* < 0.001; **, *p* < 0.01; *, *p* < 0.05.

**Fig. S6.**
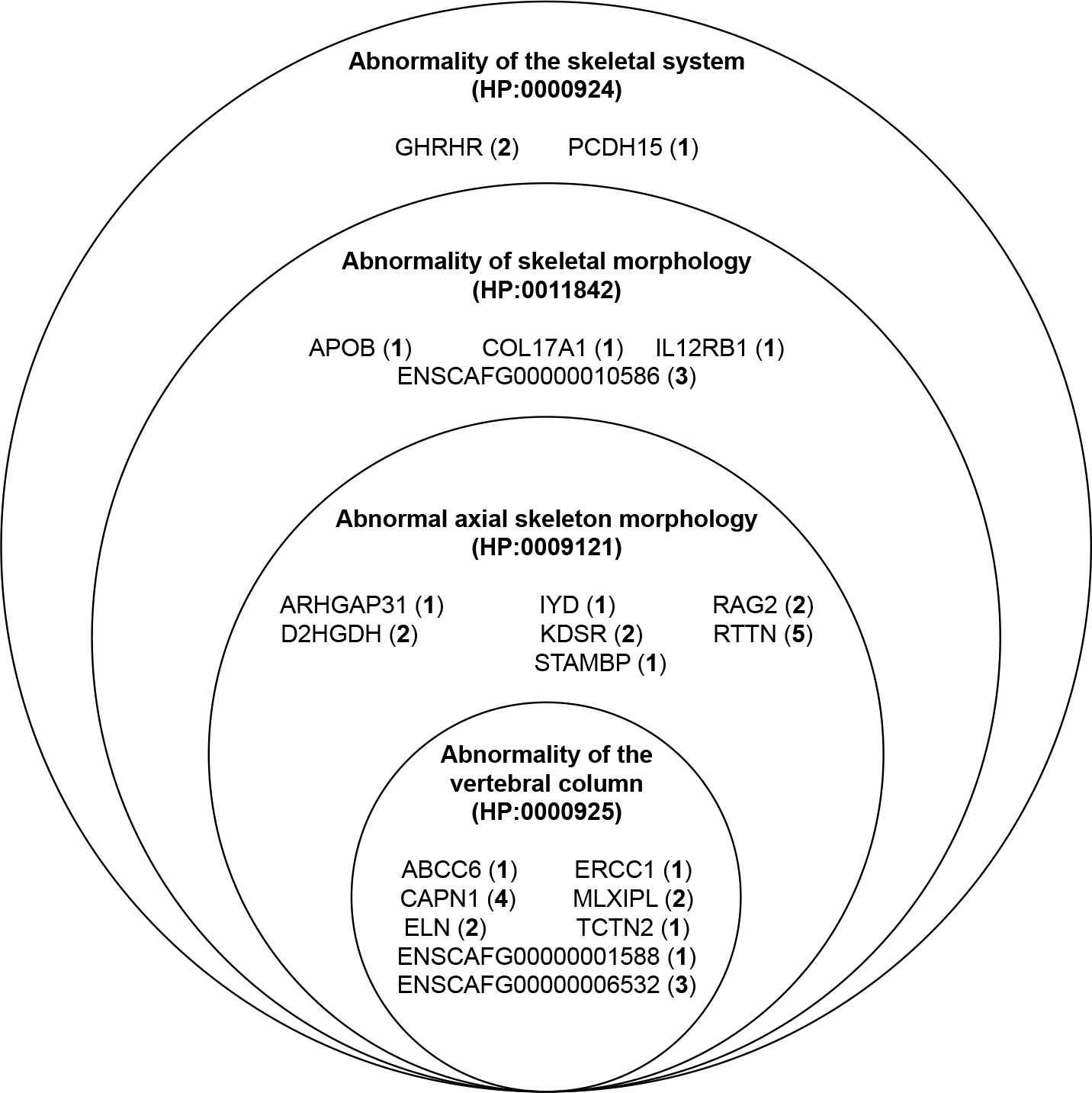
Candidate genes underlying Isle Royale phenotypes associated with HPO terms related to skeletal anatomy. Genes containing candidate mutations in ROH within affected Isle Royale wolves (F65, F75, M152, M175, F189; see Table 1 of main text) were identified. Candidate genes associated with HPO terms related to skeletal development are shown, with the number of affected individuals carrying the homozygous derived allele indicated in parentheses. HPO terms are nested, as shown.

**Table S1.** Sample information for sequences included in this study. Except where noted, analyses included all individuals listed except the African golden wolf and gray fox, which were used for polarization of alleles as ancestral or derived.

| Genome | Species | Other ID(s) | Year Sampled | Coverage (X) | Source |
| --- | --- | --- | --- | --- | --- |
| Isle Royale | *Canis lupus* | F25 | 1988 | 22.8 | This study |
| Isle Royale | *Canis lupus* | F55 | 1998 | 25.0 | This study |
| Isle Royale | *Canis lupus* | M61 | 2001 | 22.7 | This study |
| Isle Royale | *Canis lupus* | M62 | 2001 | 23.6 | This study |
| Isle Royale | *Canis lupus* | F65 | 2003 | 22.8 | This study |
| Isle Royale | *Canis lupus* | F67 | 2003 | 24.1 | This study |
| Isle Royale | *Canis lupus* | F75 | 2007 | 23.8 | This study |
| Isle Royale | *Canis lupus* | M141 | 2009 | 23.4 | This study |
| Isle Royale | *Canis lupus* | M152 | 2009 | 23.2 | This study |
| Isle Royale | *Canis lupus* | M175 | 2009 | 24.2 | This study |
| Isle Royale | *Canis lupus* | F189 | 2012 | 23.8 | This study |
| Minnesota | *Canis lupus* | RKW119 |  | 19.9 | This study |
| Minnesota | *Canis lupus* | RKW2455 |  | 24.3 | DOI:10.1101/gr.197517.115 |
| Minnesota | *Canis lupus* | RKW2515 |  | 23.4 | This study |
| Minnesota | *Canis lupus* | RKW2518 |  | 21.5 | This study |
| Minnesota | *Canis lupus* | RKW2523 |  | 22.9 | This study |
| Minnesota | *Canis lupus* | RKW2524 |  | 20.5 | This study |
| Arctic, Baffin Island | *Canis lupus* | RKW7639, CD130 |  | 49.4 | This study |
| Arctic, Ellesmere Island | *Canis lupus* | RKW7640, GF44 |  | 48.3 | This study |
| Arctic, Nunavut | *Canis lupus* | RKW7649, CB177 |  | 35.8 | This study |
| Arctic, Victoria Island | *Canis lupus* | RKW7619, CB215 |  | 31.0 | This study |
| Quebec | *Canis lupus* | 0833M |  | 25.5 | This study |
| Iran | *Canis lupus* | RKW3073 |  | 26.3 | DOI:10.1101/gr.197517.115 |
| Italy | *Canis lupus* | RKW2735, W40 |  | 12.0 | DOI:10.1101/gr.197517.115 |
| Mexico | *Canis lupus* | RKW3747, Ghost Ranch 6 |  | 23.6 | DOI:10.1101/gr.197517.115 |
| Portugal | *Canis lupus* | RKW13461, 423 |  | 24.4 | DOI:10.1101/gr.197517.115 |
| Spain | *Canis lupus* | WIB98 |  | 22.7 | DOI:10.1101/gr.197517.115 |
| Yellowstone | *Canis lupus* | RKW1547, 569F |  | 25.7 | DOI:10.1101/gr.197517.115 |
| Yellowstone | *Canis lupus* | RKW938, 302M |  | 24.1 | Provided by Rena M. Schweizer |
| Yellowstone | *Canis lupus* | RKW986, 480M |  | 18.9 | Provided by Rena M. Schweizer |
| Tibet | *Canis lupus* | TI09 |  | 24.8 | DOI:10.1371/journal.pgen.1004466 |
| Xinjiang | *Canis lupus* | XJ30 |  | 25.8 | DOI:10.1371/journal.pgen.1004466 |
| Ethiopia | *Canis simensis* |  |  | 9.1 | Provided by Thomas P. Gilbert |
| Coyote | *Canis latrans* | RKW13455, C106 |  | 24.3 | DOI:10.1101/gr.197517.115 |
| Red Wolf | *Canis rufus* | RKW701, RW179 |  | 27.2 | DOI:10.1101/gr.197517.115 |
| African Golden Wolf | *Canis aureus* |  |  | 24.8 | DOI:10.1016/j.cub.2015.06.060 |
| Gray Fox | *Urocyon cinereoargenteus* | GF41F |  | 17.0 | DOI:10.1016/j.cub.2016.02.062 |

**Table S2.**
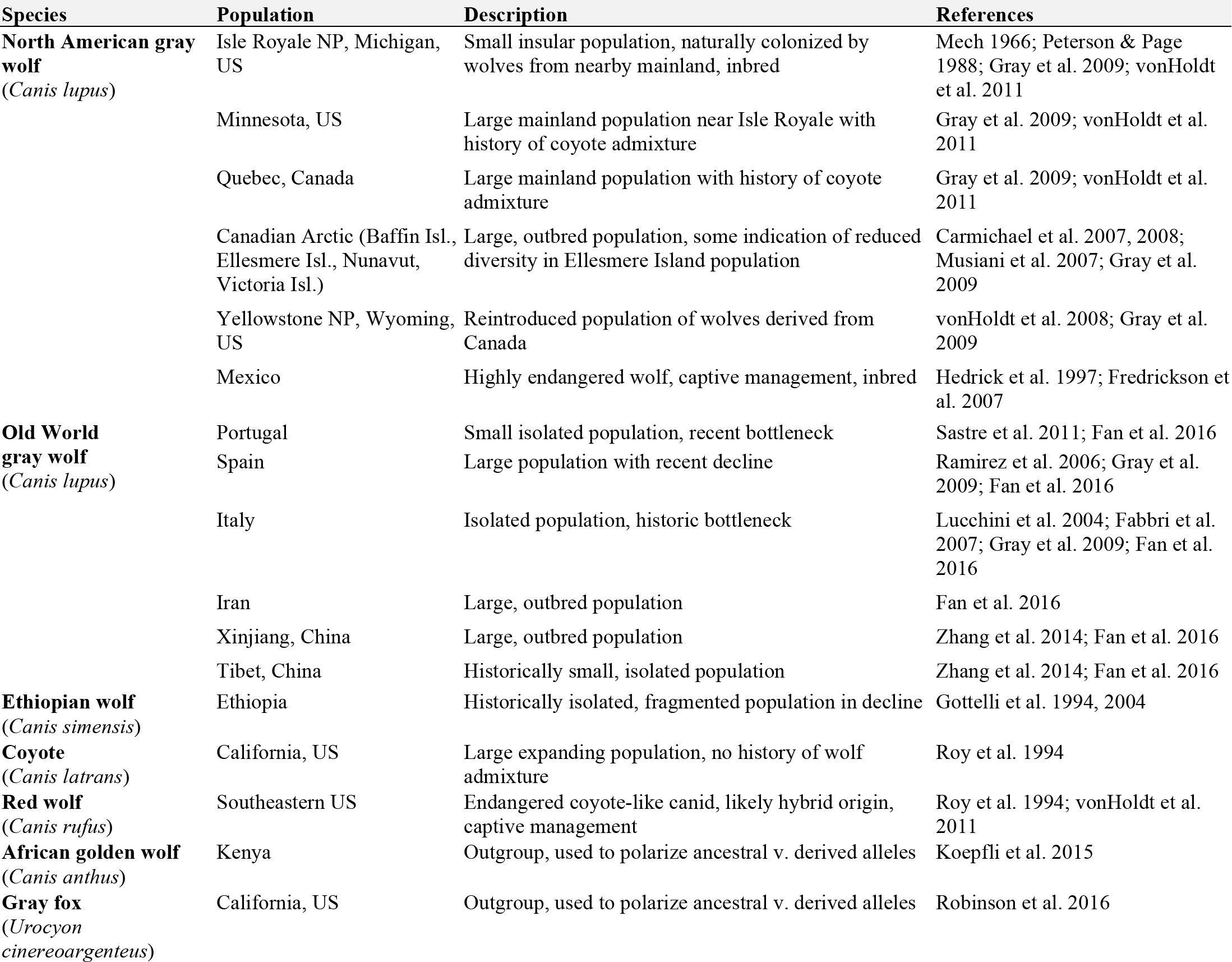
Notes and information from the literature about the demographic history of populations included in this study. Table adapted from Table 1 of vonHoldt et al. 2011.

| Species | Population | Description | References |
| --- | --- | --- | --- |
| **North American gray wolf**<br>(*Canis lupus*) | Isle Royale NP, Michigan, US | Small insular population, naturally colonized by wolves from nearby mainland, inbred | Mech 1966; Peterson & Page 1988; Gray et al. 2009; vonHoldt et al. 2011 |
|  | Minnesota, US | Large mainland population near Isle Royale with history of coyote admixture | Gray et al. 2009; vonHoldt et al. 2011 |
|  | Quebec, Canada | Large mainland population with history of coyote admixture | Gray et al. 2009; vonHoldt et al. 2011 |
|  | Canadian Arctic (Baffin Isl., Ellesmere Isl., Nunavut, Victoria Isl.) | Large, outbred population, some indication of reduced diversity in Ellesmere Island population | Carmichael et al. 2007, 2008; Musiani et al. 2007; Gray et al. 2009 |
|  | Yellowstone NP, Wyoming, US | Reintroduced population of wolves derived from Canada | vonHoldt et al. 2008; Gray et al. 2009 |
|  | Mexico | Highly endangered wolf, captive management, inbred | Hedrick et al. 1997; Fredrickson et al. 2007 |
| **Old World gray wolf**<br>(*Canis lupus*) | Portugal | Small isolated population, recent bottleneck | Sastre et al. 2011; Fan et al. 2016 |
|  | Spain | Large population with recent decline | Ramirez et al. 2006; Gray et al. 2009; Fan et al. 2016 |
|  | Italy | Isolated population, historic bottleneck | Lucchini et al. 2004; Fabbri et al. 2007; Gray et al. 2009; Fan et al. 2016 |
|  | Iran | Large, outbred population | Fan et al. 2016 |
|  | Xinjiang, China | Large, outbred population | Zhang et al. 2014; Fan et al. 2016 |
|  | Tibet, China | Historically small, isolated population | Zhang et al. 2014; Fan et al. 2016 |
| **Ethiopian wolf**<br>(*Canis simensis*) | Ethiopia | Historically isolated, fragmented population in decline | Gottelli et al. 1994, 2004 |
| **Coyote**<br>(*Canis latrans*) | California, US | Large expanding population, no history of wolf admixture | Roy et al. 1994 |
| **Red wolf**<br>(*Canis rufus*) | Southeastern US | Endangered coyote-like canid, likely hybrid origin, captive management | Roy et al. 1994; vonHoldt et al. 2011 |
| **African golden wolf**<br>(*Canis anthus*) | Kenya | Outgroup, used to polarize ancestral v. derived alleles | Koepfli et al. 2015 |
| **Gray fox**<br>(*Urocyon cinereoargenteus*) | California, US | Outgroup, used to polarize ancestral v. derived alleles | Robinson et al. 2016 |

**Table S3.**
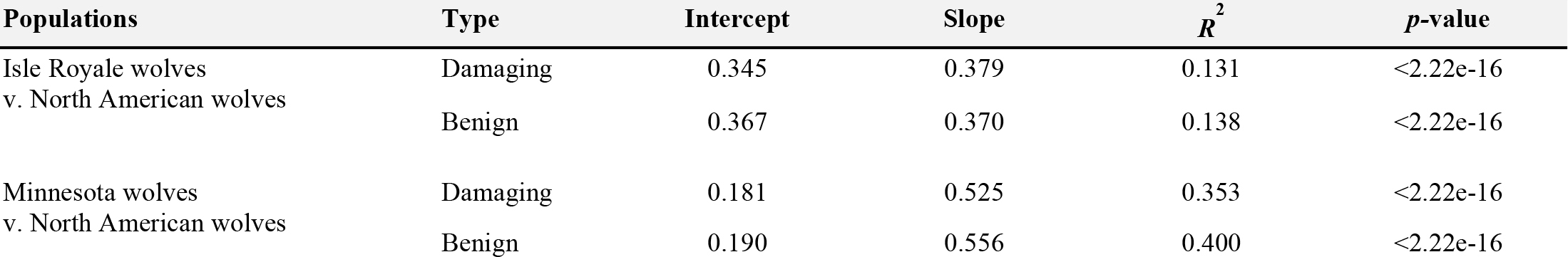
Parameters of linear regression models for two-dimensional allele frequency spectra. Spectra are shown in main text (Fig. 4 D-G).

